# Outer Kinetochore Proteins form Linear Elements to Regulate Vesicle Transport

**DOI:** 10.1101/2025.04.25.650656

**Authors:** Shane J. Kowaleski, Alexis Bridgewater, Cody Saraceno, Miranda Dudek, Federico Pelisch, Joshua N. Bembenek

## Abstract

During cell division, several key regulators of chromosome segregation play additional roles during vesicle trafficking required for cytokinesis. During anaphase I in *C. elegans* oocytes, chromosome segregation is coordinated with vesicle trafficking to support polar body extrusion and exocytosis of extracellular matrix material. Prior to anaphase, numerous outer kinetochore proteins localize to mysterious “linear element” structures throughout the cortex in addition to chromosomes, which has been observed in oocytes of multiple species. Linear elements initially form as puncta just before nuclear envelope breakdown and rapidly assemble into larger elongated structures. As linear elements grow, they form large clusters with secretory vesicles, initiating an elaborate transport mechanism that distributes vesicles throughout the cortex by anaphase I. Linear elements dynamically interact with microtubules and endoplasmic reticulum during this process. Microtubules are required for linear element assembly, motility, and clustering with vesicles. Knockdown of a plus end microtubule binding kinetochore component also inhibits linear element growth and vesicle clustering, but not the motility of linear element puncta. Depletion of several outer kinetochore proteins causes defects in extracellular matrix formation. Therefore, linear elements facilitate the microtubule-dependent transport of vesicles for their proper distribution in the cortex. We hypothesize that outer kinetochore complexes coordinate movements of chromosomes and cytoplasmic membranes to enhance the fidelity of cell division.

## Introduction

The kinetochore is a mechanosensitive microtubule coupling device and signaling platform required for chromosome segregation during cell division^1,2^. Structurally, kinetochores are multilayered assemblies at centromeres, with an outer layer that captures microtubules^3,4^. A critical signaling pathway known as the spindle assembly checkpoint monitors microtubule attachment and tension at kinetochores and inhibits anaphase onset^5,6^. Once proper chromosome alignment is achieved during metaphase, the checkpoint is silenced, allowing the activation of separase^7^. Separase is a protease that cleaves cohesin and allows the poleward movement of chromosomes at anaphase onset^8^.

Separase has multiple functions during anaphase, including the regulation of spindle dynamics^9–11^, CDK activity^8,12^ and centriole licensing^13,14^. Separase activity is also implicated in the regulation of cytokinesis in several systems^15–17^. We discovered that separase is required for cytokinesis by promoting exocytosis during anaphase^18^. In general, it is thought that cells shut down trafficking during early stages of cell division and resume exocytosis during anaphase to expand the plasma membrane during cytokinesis^19–22^. A major exocytic pathway involves RAB-11 dependent vesicle trafficking from a centrosomal pool of membrane to the plasma membrane^23–26^. The regulation of RAB-11 vesicle trafficking by separase allows the cell to stimulate exocytosis in coordination with the onset of chromosome separation at anaphase onset.

During oocyte meiosis I in *C. elegans*, homologous chromosomes are segregated, and one set is partitioned into a small polar body during a highly asymmetric cytokinesis^27^. These events occur during egg activation, a collection of events triggered in oocytes after fertilization^28^. Another major egg activation event is the release of cargo through cortical granule exocytosis to modify the extracellular matrix, which provides physical and osmotic protection to the developing embryo^29^ as well as forms the polyspermy barrier in several species^30^. Numerous cell cycle genes have been identified that affect extracellular matrix formation, implicating them in secretion^31,32^. Importantly, both separase and RAB-11 localize to cortical granules and are required for their exocytosis in anaphase I^33,34^. This finding implies that the spindle assembly checkpoint pathway, at least through regulation of separase activity, also controls the timing of exocytosis in anaphase. The full extent of cell cycle regulation of vesicle trafficking is still largely unexplored.

Separase localizes to kinetochores and linear elements at early stages of division and only moves to sites of action at anaphase onset^33,35^. Numerous outer kinetochore proteins have also been shown to localize to cytoplasmic linear element structures in *C. elegans*^36–45^, *Drosophila*^46–48^ and bovine oocytes^49^. While these structures are not well characterized, their colocalization with separase in the cortex implies a potential indirect role in regulating vesicle trafficking. Several interesting observations suggest outer kinetochore components might have membrane related functions. The ROD-ZWILCH-ZW10-Spindly (RZZ-S) complex has structural similarity with vesicle coat proteins^50,51^. ZW10 binds to Syntaxin 18 in an endoplasmic reticulum (ER) tethering complex to regulate ER-Golgi trafficking^52,53^. Multiple kinetochore proteins including Spindly^54^, CENP-E^55,56^ and CENP-F^57^ are farnesylated, a posttranslational modification that can facilitate membrane association^58^. ZWINT interacts with Rab3c and SNAP25 and is implicated in vesicle exocytosis^59,60^. Additionally, the SNARE protein SNAP29 recruits KNL-1 to the kinetochore^61^ and copurifies with Ska1^62^. Therefore, kinetochore proteins may regulate various aspects of membrane trafficking.

Here, we demonstrate that linear elements form in *C. elegans* oocytes just before nuclear envelope breakdown (NEBD) from smaller puncta, as previously observed in *Drosophila* oocytes^46^. We show that during their formation, linear elements form large clusters with cortical granule vesicles during a complex vesicle transport process that results in vesicle distribution throughout the cortex. Linear elements continuously interact with endoplasmic reticulum and microtubules during their formation. We investigate the function of linear elements during vesicle trafficking and uncover a novel role for outer kinetochore proteins at membranous organelles that occurs concurrently with their well-known function on chromosomes.

## Results

### Linear Element Dynamics During Meiosis I

We recently characterized separase (SEP-1 in *C. elegans*) localization to kinetochores and linear elements and determined that it moves to sites of action during anaphase of oocyte meiosis I^35^. While observing separase localization, we observed that linear elements form just before nuclear envelope breakdown, which marks the start of prometaphase I (N=9, Figure 1A-1D). We generated a strain expressing endogenously tagged SEP-1::mScarlet and transgenic CZW-1::GFP, the ZW10 homologue, to observe linear element dynamics. Initially, small CZW-1::GFP puncta form and rapidly assemble into longer structures (Figure 1B, 1E and 1F), with SEP-1::mScarlet recruited slightly later (N=4, Figure 1C and 1F). CZW-1::GFP is observed on chromosomes before linear elements are observed, and separase is only recruited to chromosomes after linear elements are fully formed (N=9, Figure 1D, 1F). Numerous outer kinetochore components and related cell cycle regulators have been reported to associate with linear elements, as summarized in Table 1. We verified numerous outer kinetochore complex components, spindle checkpoint regulators and several microtubule-associated proteins all show prominent localization to linear elements, but not the inner kinetochore protein CENP-C (Figure S1). Linear elements persist in the cortex until mid-anaphase I, when they disappear just before the time that SEP-1 relocalizes to cortical granules (N=8, Figure 1G). Therefore, linear elements form in the cortical cytoplasm prior to nuclear envelope breakdown and persist until anaphase when separase moves to vesicles.

**Figure 1.**
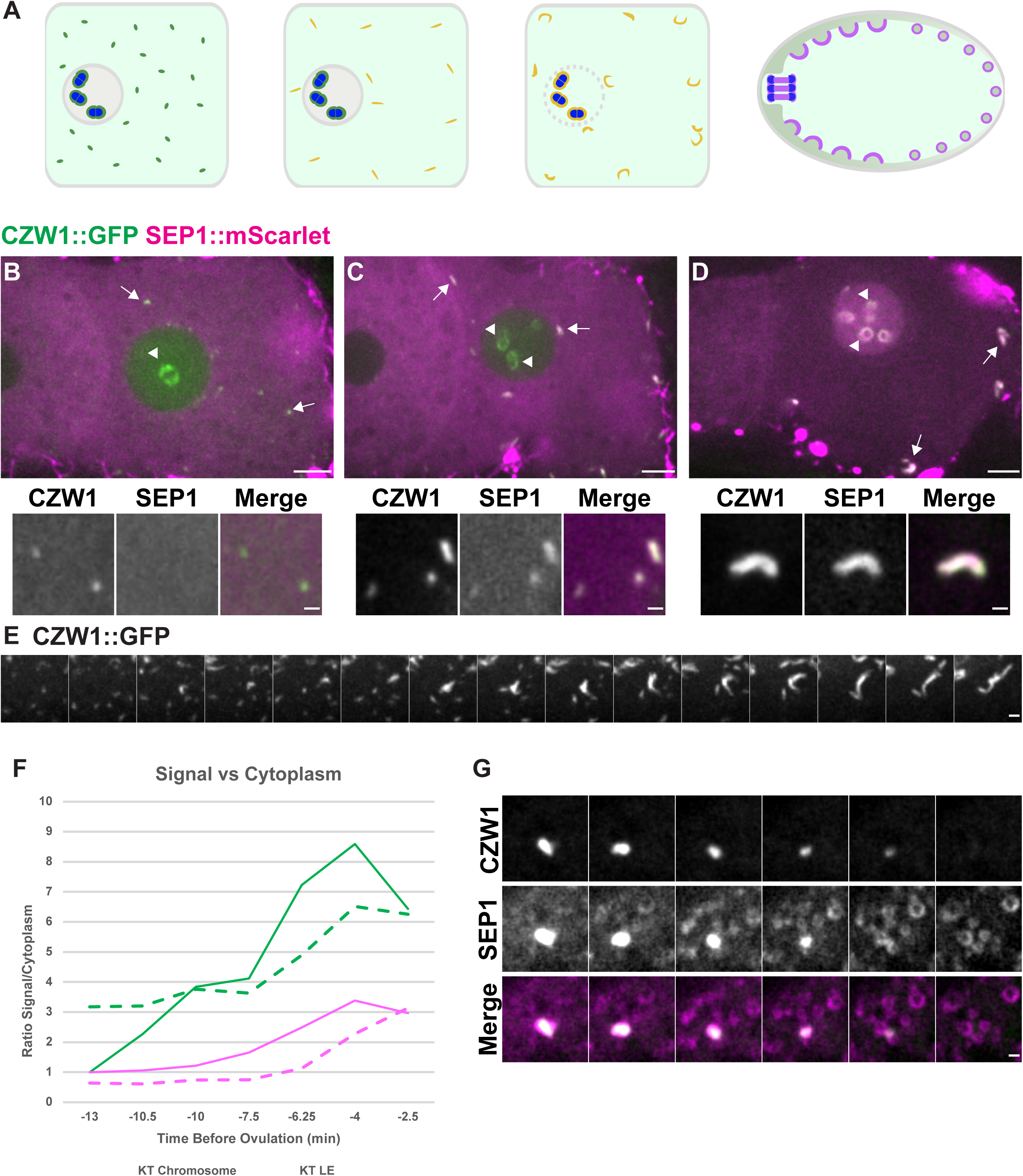
Linear Element Dynamics During Meiosis. **I.** (A) Diagrams of *C. elegans* oocyte meiosis I showing linear element formation. ZW10^czw-1^ (green, Separase^sep-1^ is magenta, colocalization is indicated by yellow) initially forms small puncta, which rapidly assemble into linear elements (middle images). Separase associates with linear elements as they grow and associates with chromosomes (blue) at NEBD. Linear elements disappear at anaphase onset (right image), while separase moves to vesicles and promotes exocytosis. (B) CZW-1::GFP (green) initially appears on small puncta (arrows) dispersed throughout the cytoplasm and is localized to chromosomes (arrowhead), while SEP-1::mScarlet (magenta) is diffuse in the cytoplasm. Upper images in B and C are maximum intensity projections, insets are single plane. Scale bar for upper image is 5 µm, inset is 1 µm. (C) SEP-1::mScarlet colocalizes with CZW-1::GFP at linear elements (arrows) when they assemble into intermediate sized structures but is not yet on chromosomes (arrowheads). (D) By NEBD, SEP-1::mScarlet colocalizes with CZW-1::GFP on fully assembled linear elements and chromosomes. Images are single plane, scale bar for upper image is 5 µm, inset is 1 µm. (E) Montage showing CZW-1::GFP puncta that rapidly assemble into larger structures. Images are maximum intensity projections, acquired every 10 seconds. Scale Bar is 1 µm. (F) Quantification of CZW-1::GFP (green) and SEP-1::mScarlet (magenta) signal on linear elements (solid lines) and chromosomes (dashed lines) over time in a representative embryo showing CZW-1::GFP accumulates on linear elements before SEP-1::mScarlet. CZW-1::GFP is already on chromosomes before it appears on linear elements, while separase accumulates on linear elements before it enters the nucleus and associates with chromosomes. (G) Montage showing CZW-1::GFP (green) and SEP-1::mScarlet (magenta) colocalize until seconds after anaphase onset, where CZW-1::GFP disperses and SEP-1 accumulates on cortical granules. Single plane images acquired every 10 seconds. Scale Bar: 1 µm.

**Table 1.**
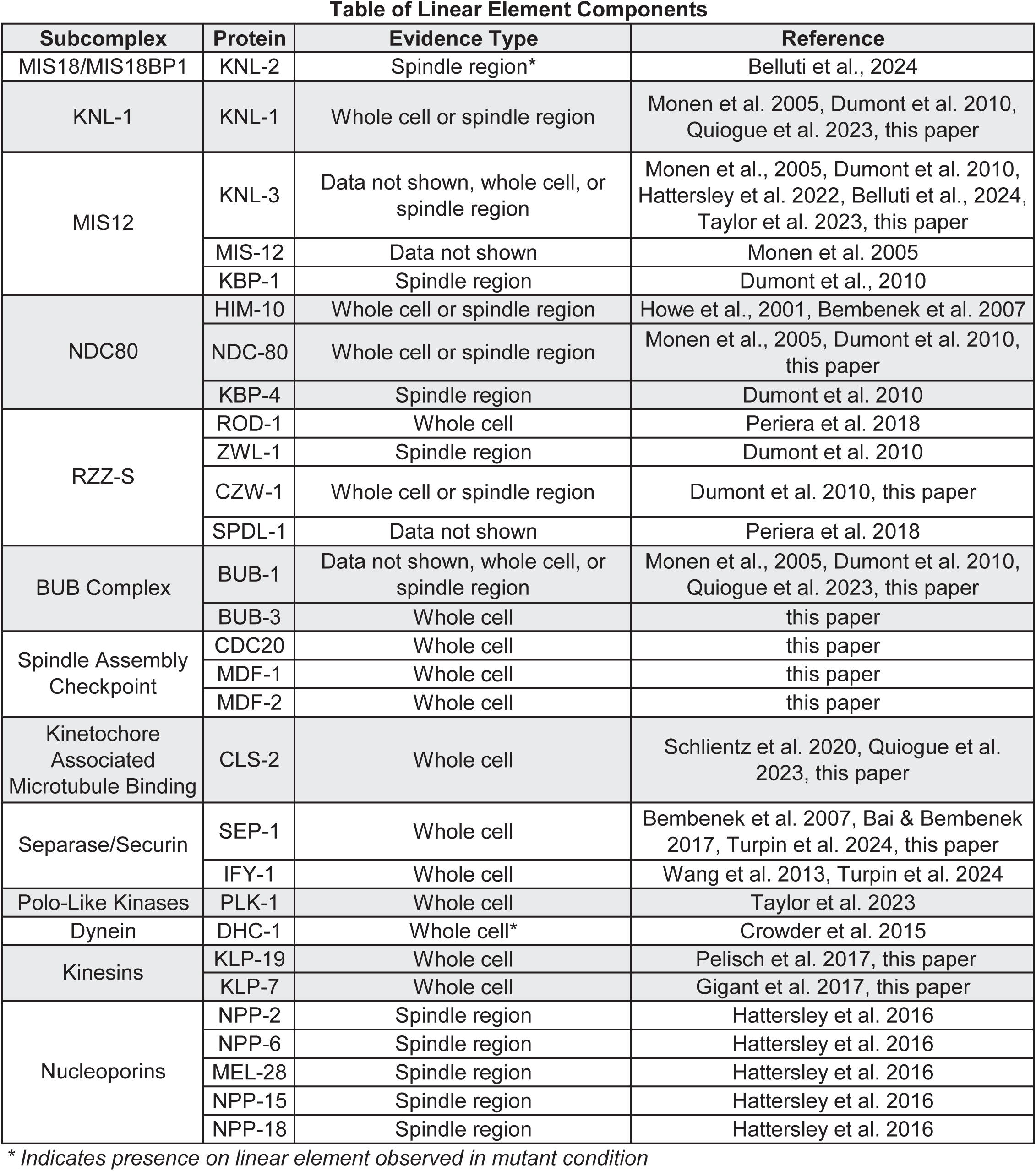

### Linear Elements Assemble into Clusters with Cortical Granules before NEBD

While investigating the dynamics of separase localization to vesicles, we made a striking observation that linear elements interact with vesicles at NEBD. We generated a strain expressing CZW-1::GFP and CPG-2::mCherry, a cargo protein of cortical granules, to document linear element and vesicle dynamics. When linear element puncta first form, they are not closely associated with cortical granules (N=5, Figure 2A). As linear elements assemble into larger structures, they dynamically interact with cortical granules and increasingly associate with them (N=6, Figure 2B). As linear elements reach their full length, they aggregate with clusters of cortical granules, which is mostly finished by nuclear envelope breakdown (N=7, Figure 2C, 2D). Therefore, linear elements assemble into large clusters with cortical granule vesicles when they form just prior to NEBD, suggesting that linear elements might control vesicle movements.

**Figure 2.**
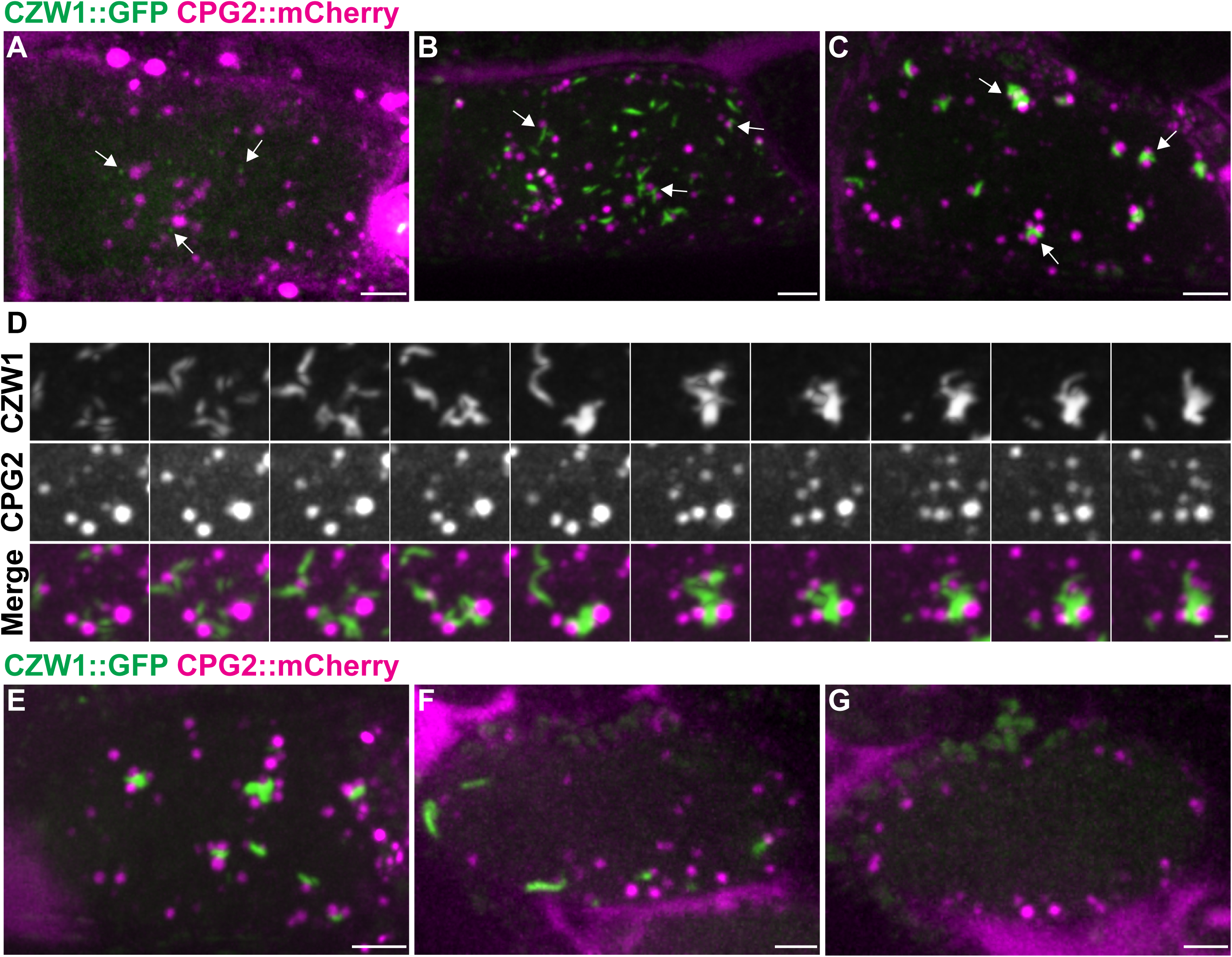
Linear Elements Associate with Cortical Granules. (A) CZW-1::GFP (green) puncta initially appear in the cortical cytoplasm away from cortical granules marked by CPG-2::mCherry (magenta). Images in A-C, E-G are maximum intensity projections, scale bars are 5 µm. (B) As linear elements assemble in the cortex (green), they move and begin to associate with cortical granules (magenta). (C) Fully assembled linear elements (green) associate with large clusters of cortical granules (magenta) in the cortex, which is completed before NEBD. (D) Montage showing CZW-1::GFP (green) assembling into a linear element and gathering CPG-2::mCherry labeled cortical granules (magenta) into clusters. Images are maximum intensity projections every 30 seconds, scale bar is 1 µm. (E) Clusters of cortical granules (magenta) and linear elements (green) remain associated through ovulation. (F) During cytoplasmic streaming, cortical granules (magenta) move in the cortex and disassociate from linear elements, which remain relatively stationary. (G) Just before exocytosis in anaphase I, cortical granules (magenta) are distributed throughout the cortex while linear elements disappear.

Given the obvious clustering of vesicles and linear elements at NEBD, we sought to place this event within the global dynamics of vesicle movements during oocyte meiosis I. Previously, cortical granule vesicles were observed in close association with the ER, which also undergoes dramatic reorganization^33,63^. Therefore, we generated a strain expressing SP12::GFP to visualize the ER, and CPG-2::mCherry to mark cortical granules. During the extensive prophase I arrest that precedes NEBD, oocytes grow in size and stockpile material necessary for embryonic development. During this “production phase”, cortical granules and ER are relatively evenly distributed throughout the cytoplasm (N=37, Figure S2B). Over a 20-30 minute period prior to NEBD, the oocyte nucleus migrates toward the distal oocyte membrane^64^. After this period, cortical granules are cortically displaced and the ER is consolidated into thick sheets that are also biased toward the cortex (N=27, Figure S2C). The ER and cortical granules remain cortically displaced through NEBD (N=19, Figure S2D). Around the time of fertilization, which occurs during prometaphase I in the oocyte, a dramatic period of cytoplasmic streaming occurs^65^. During streaming, the ER becomes more diffuse but still cortically displaced and rapidly moves with cortical granules (N=5, Figure S2E). Finally, vesicles undergo exocytosis in mid anaphase (N=5, Figure S2F). Therefore, linear elements cluster with cortical granules in the middle of their journey from biosynthesis to fusion with the plasma membrane.

We observed the dynamics of linear elements relative to cortical granules during the different stages of vesicle distribution. Linear elements form several minutes prior to NEBD, when the ER and cortical granules are already cortically displaced (Figure 2A, S1C) and remain associated with them through ovulation (N=5, Figure 2E). Around fertilization, cortical granules begin to dissociate from linear elements and cytoplasmic streaming ensues (N=4, Figure 2F). During cytoplasmic streaming, linear elements are relatively stationary in the cortex while cortical granules move quickly. Linear elements are observed in the cortex until mid-anaphase I, when they disappear at the time that separase moves onto vesicles as we recently described (N=4, Figure 2G)^35^. Therefore, cortical granules undergo a complex transport process to become distributed throughout the cortex by anaphase I and closely interact with linear elements transiently just prior to NEBD.

### Linear Elements Associate with the ER

Since cortical granules interact with the ER throughout meiosis I, we sought to investigate whether linear elements also interact with the ER. We generated a strain containing the outer kinetochore protein KNL-1 endogenously tagged with TagRFP to label linear elements and exogenous SP12::GFP to label the ER. Linear element puncta associate with ER at the time of their formation (N=4, Figure 3A). This association is maintained during their assembly, with intermediate sized linear elements continuously interacting with the ER (N=7, Figure 3B). When linear elements reach their full size, they associate with domains of ER throughout the cytoplasm and adjacent to the nucleus (N=10, Figure 3C), which is maintained after ovulation around the time of fertilization (N=10, Figure 3D). During cytoplasmic streaming in prometaphase I, linear elements remain static while the ER flows in the cortex (N=5, Figure 3E). In mid-anaphase I, linear elements disappear while cortical ER remains distributed throughout the cortex (N=2, Figure 3F). Therefore, linear elements interact with ER throughout their formation and then remain static in the cortex during cytoplasmic streaming as ER and vesicles move rapidly.

**Figure 3.**
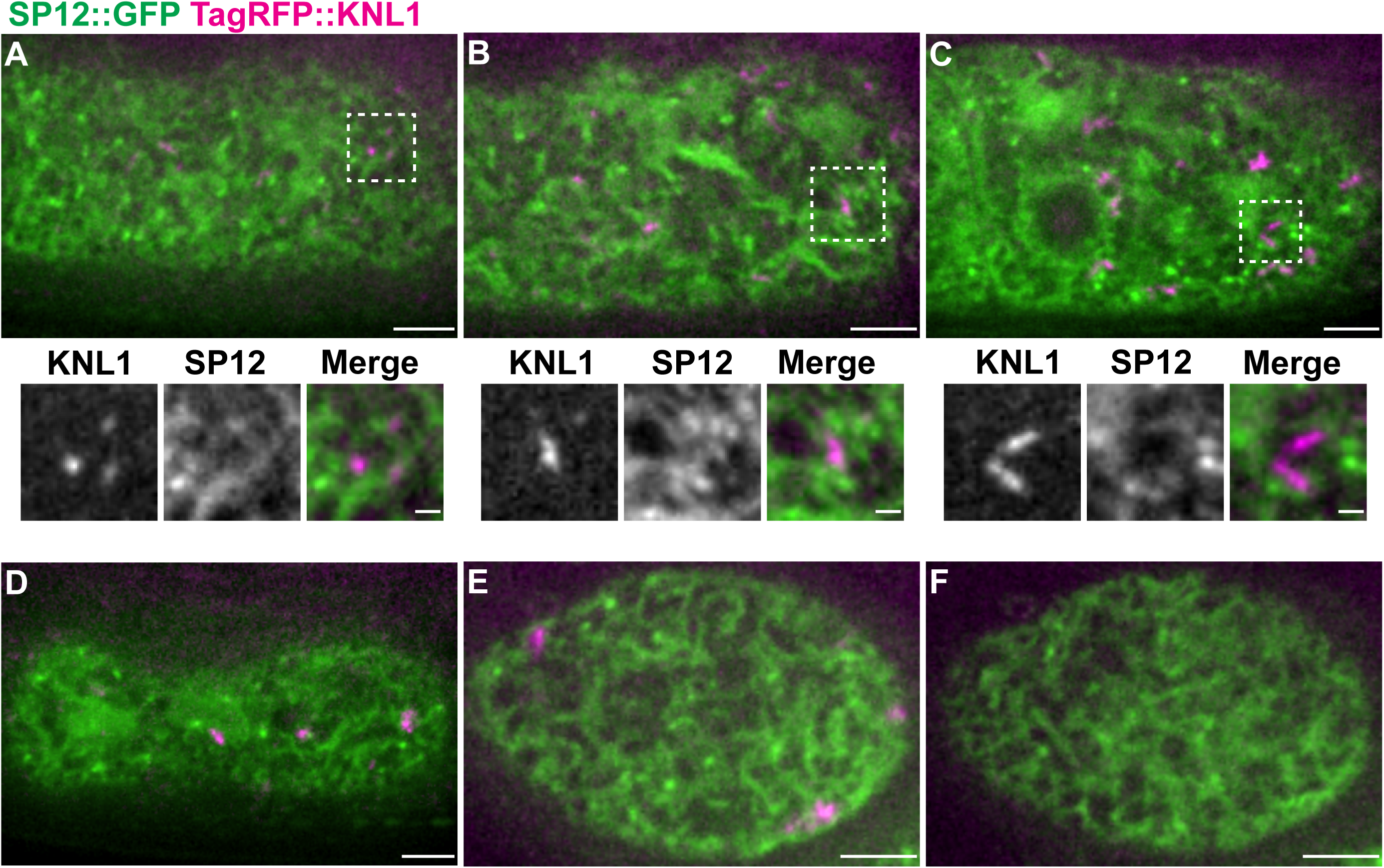
Linear Elements Associate with the Endoplasmic Reticulum. (A) Cortical planes showing TagRFP::KNL-1 (magenta) initially forms small puncta adjacent to domains of ER marked by SP12::GFP (green). A-F are single plane images, scale bar for whole cell images in A-F is 5 µm, insets are 1 µm. (B) Linear elements (magenta) remain embedded within the cortical ER network as they move together and assemble into larger structures. (C) Fully assembled linear elements (magenta) are surrounded by domains of cortical ER (green) just before nuclear envelope breakdown. (D) Linear elements (magenta) remain associated with the cortical ER (green) during ovulation. (E) During cytoplasmic streaming, the ER rapidly flows in the cortex, while linear elements are more static and adjacent to ER. (F) Cortical ER morphology after linear element disappearance in anaphase I.

### Linear Elements Interact with the Cortical Microtubule Network

The critical function of the kinetochore is to mediate spindle microtubule attachment to chromosomes. Therefore, we wanted to investigate the relationship between the linear elements and the cortical microtubule cytoskeleton. We generated a strain expressing endogenously tagged TagRFP::KNL-1 and GFP::tubulin and imaged oocytes during linear element formation just before NEBD. We observed that linear element puncta overlap or appear adjacent to microtubules when they first form (N=9, Figure 4A). As linear elements assemble, they accumulate microtubule bundles within an extended microtubule network that in some cases appears to connect between linear element foci (N=6, Figure 4B). When linear elements are fully elongated, we observe asymmetric accumulations of microtubules at different points along their length (N=8, Figure 4C). Linear elements remain affiliated with microtubules through fertilization (N=3, Figure 4D), dynamically interact with the microtubules during streaming (N=7, Figure 4E), up until their disappearance in mid-anaphase I, when microtubule distribution is more even (N=5, Figure 4F). Therefore, linear elements constantly interact with the dynamic cortical microtubule network.

**Figure 4.**
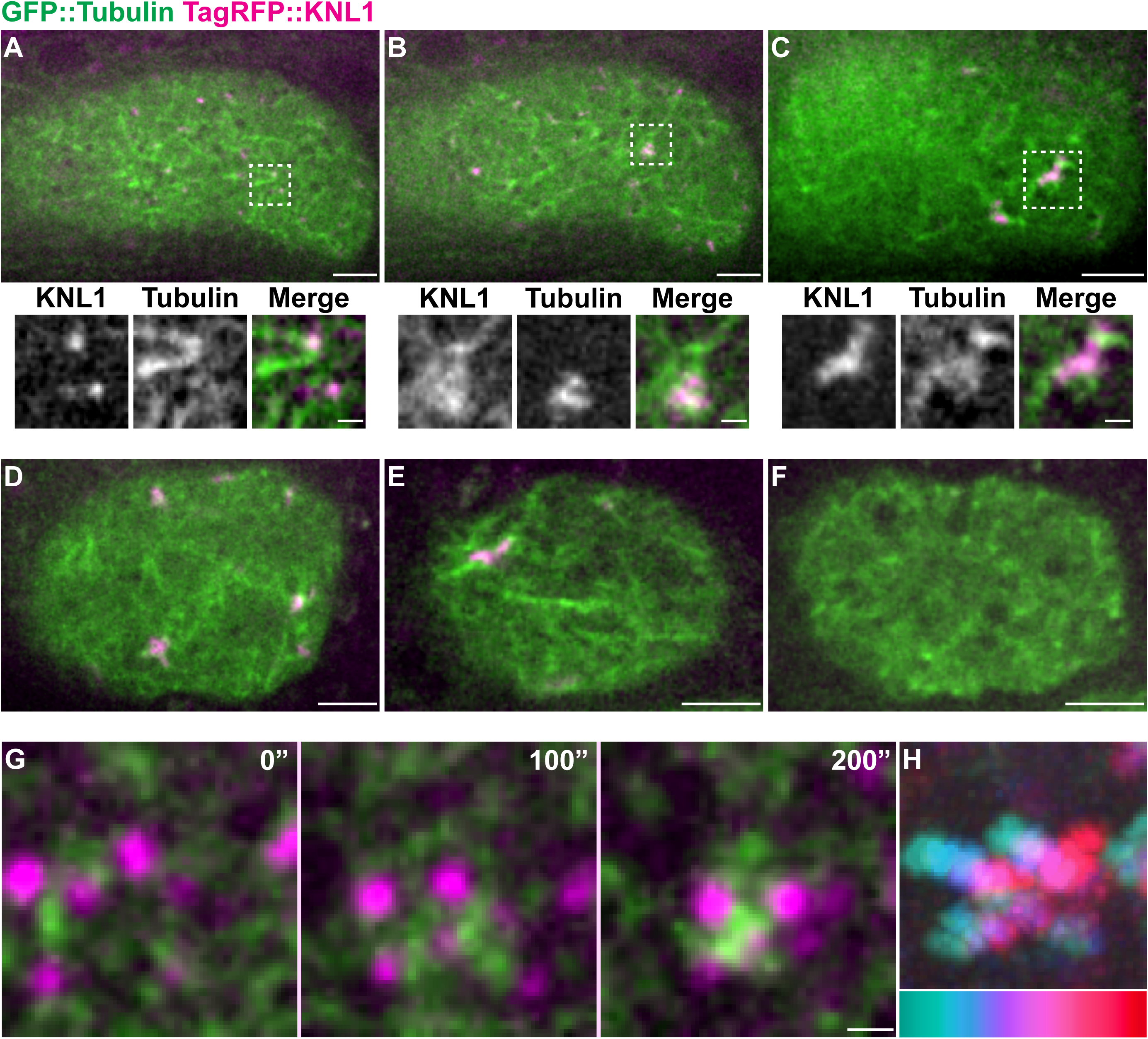
Linear Elements Interact with the Cortical Microtubule Network. (A) Linear element puncta marked by TagRFP::KNL-1 (magenta) appear within the cortical microtubule network (green). A-F are single plane images, whole cortex view scale bars are 5 µm, insets are 1 µm. (B) As linear elements (magenta) assemble, they constantly interact with cortical microtubules (green), which reorganize and assemble into bundles. (C) Fully assembled linear elements (magenta) continue to interact with cortical microtubules (green), with accumulations of microtubules in different regions along the linear element. (D) Linear elements (magenta) remain associated with microtubules (green) during ovulation. (E) During cytoplasmic streaming, microtubules (green) flow continuously in the cortex and dynamically interact with linear elements, which are relatively static. (F) Cortical microtubule network morphology during anaphase I after linear elements disappear. (G) Cortical granules marked by CPG-2::mCherry (magenta) are surrounded by cortical microtubules (green). Microtubules (green) accumulate as cortical granules gather into clusters (second and third panels). 100 seconds separate panels. Scale Bar: 1 µm. (H) Color coded temporal projection of vesicle clustering, indicating inward movement of vesicles. Time interval is 5 seconds.

We also investigated the interaction between cortical granules and microtubules using a strain expressing GFP::tubulin and CPG-2::mCherry. Cortical granules are within the cortical microtubule network prior to nuclear envelope breakdown. As the oocyte gets closer to nuclear envelope breakdown, around the time when linear elements form, cortical granules can be observed packing into clusters that contain a high density of microtubules (N=8, Figure 4G, 4H). Collectively, these results indicate that linear elements form in contact with microtubules and the ER, and as they grow, they interact with cortical granule vesicles and form large clusters. These results suggest that linear elements may regulate vesicle transport in the oocyte.

### Kinetochore Proteins and Microtubules are Required for Vesicle Clustering

We next sought to test the hypothesis that linear elements are required for microtubule dependent vesicle transport. We investigated vesicle movements just before NEBD, using a line expressing CZW-1::GFP and CPG-2::mCherry. To test the role of microtubules in this process, we performed *tba-2* feeding RNAi for 20-28 hours. Unlike control animals, where linear elements form quickly and cluster with vesicles before NEBD (N=9/9, Figure 5A-C, M, top row), linear elements in *tba-2(RNAi)* oocytes remained smaller and distributed throughout the cortex through ovulation (N=12/14, Figure 5D-F). Previously, linear element structures have been observed to grow larger after microtubule disruptions^41,49^. To resolve this apparent discrepancy, we analyzed linear elements in APC/C-inactivated embryos and found that linear elements can eventually grow larger in long-term arrested embryos depleted of microtubules (Figure S3). Linear elements did not cluster with vesicles and were relatively immobile after *tba-2(RNAi)* compared with controls (Figure 5K, 5J). Therefore, microtubules are required for efficient linear element assembly and vesicle clustering.

**Figure 5.**
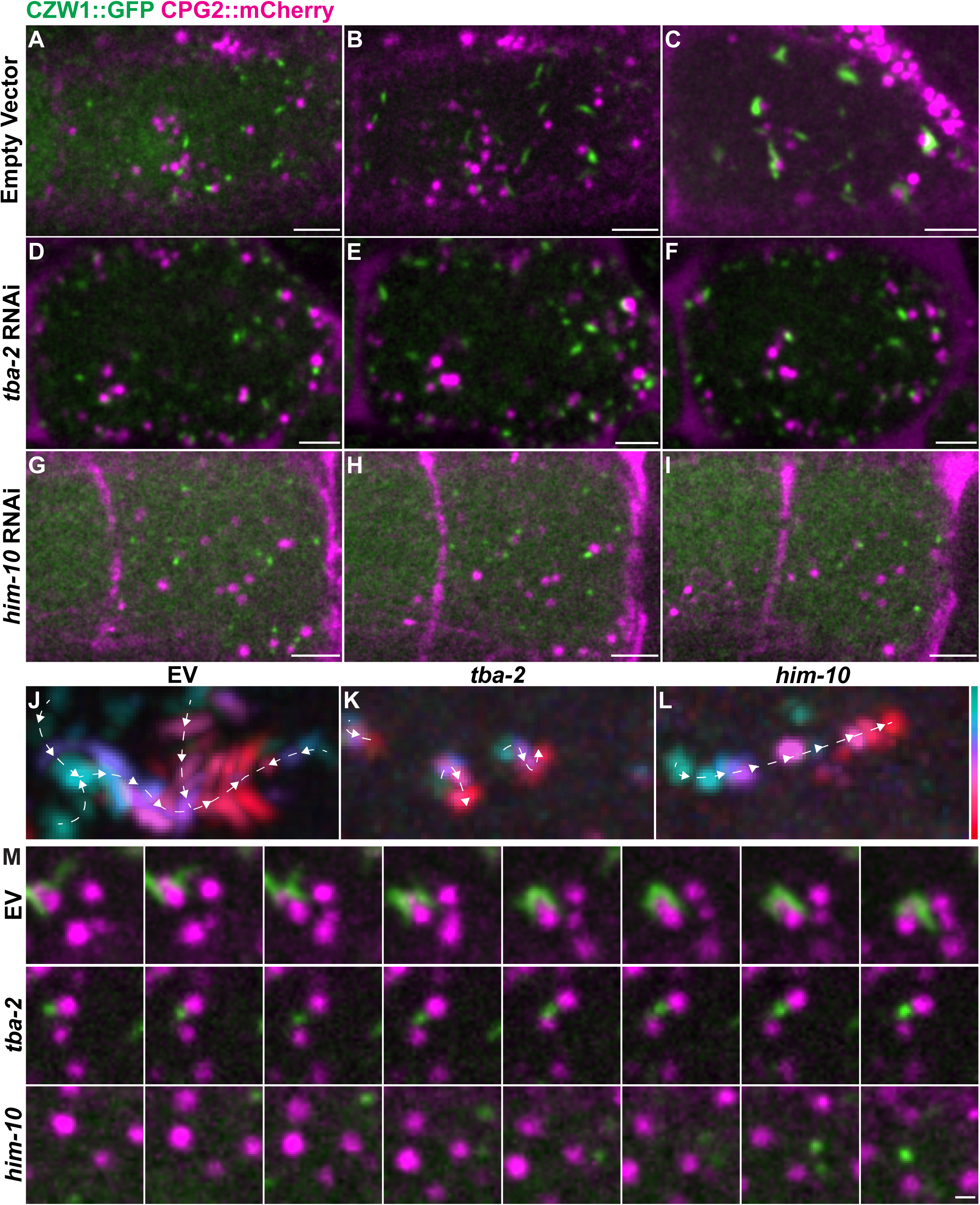
Microtubules and Linear Elements are Required for Vesicle Movement. (A-C) Cortical planes showing CZW-1::GFP forming linear elements (green) and clustering with CPG-2::mCherry labeled cortical granules (magenta). Small puncta (A) assemble larger (B) and become full length (C) in control oocytes just before NEBD. Scale bars are 5 µm. (D-F) Depletion of microtubules in *tba-2(RNAi)* oocytes severely inhibits linear element (green) assembly and motility. In addition, linear elements do not form clusters with cortical granules (magenta) at an early (D) middle (E) or late (F) timepoints. Scale bars are 5 µm. (G-I) In *him-10(RNAi)* oocytes, small linear element puncta (green) do not assemble into larger structures at early (G), middle (H), and late (I) timepoints, but are still motile in the cortex. Furthermore, cortical granules (magenta) do not cluster. Scale bars are 5 µm. (J-L) Color coded temporal projections of linear elements in control (J), *tba-2(RNAi)* (K), and *him-10(RNAi)* (L) treated oocytes. Linear elements move extensively in control and *him-10(RNAi)*, but are stationary after *tba-2(RNAi)*. Time interval is 5 seconds. (M) Montage of linear element (green) and cortical granule (magenta) cluster formation in control (top), *tba-2(RNAi)* (middle), and *him-10(RNAi)* (bottom) oocytes. Vesicle clustering occurs in control, but not *tba-2(RNAi)* or *him-10(RNAi)*. Time intervals are 10 seconds, scale bars are 1 µm.

Additionally, cortical granule distribution appeared abnormal prior to linear element formation, indicating an earlier defect in their positioning. To further characterize the role of microtubules in vesicle dynamics, we examined ER and cortical granule distribution after *tba-2(RNAi)* at different stages of meiosis I. In prophase I arrested *tba-2(RNAi)* oocytes, vesicles remain closer together and the ER is less dispersed throughout the cytoplasm than control (N=5, Figure S2G). Around NEBD, the ER and vesicles are not as cortically displaced in *tba-2(RNAi)* oocytes (N=15, Figure S2H, S2I) and they remain displaced from the cortex during the time when streaming should happen through exocytosis in anaphase I (N=4, Figure S2J, S2K). Critically, we observe vesicles that remain trapped in the cytoplasm after the majority undergo exocytosis anaphase I (N=10/11, Figure S2K). Therefore, microtubule-dependent transport is required for proper cortical granule transport and distribution in the cortex, which is a prerequisite for exocytosis.

To determine whether kinetochore proteins within linear elements are required for vesicle movements, we depleted the Nuf2 homologue, *him-10*, an NDC80 complex component involved in plus end microtubule binding. After 41-46 hours of feeding *him-10* (RNAi), we observed linear element formation and vesicle clustering at NEBD. In *him-10(RNAi)* oocytes, linear element puncta are severely inhibited from assembling into larger structures and do not form clusters with cortical granules, which remain dispersed in the cortex until ovulation (N=10/12, Figure 5G-I, M, bottom row). Interestingly, linear element puncta in *him-10(RNAi)* oocytes are still motile and move dynamically in the cortex (Figure 5L). Therefore, Nuf2*^him-^*^10^ is required for linear element assembly, but is not required for their motility. Finally, we performed a small-scale RNAi screen using Nile Blue plates to detect permeable embryos, an indication that eggshell formation is disrupted. Depletion of numerous outer kinetochore proteins leads to production of dead embryos, a small but significant fraction that show Nile Blue staining (Figure S4). Therefore, these results indicate that linear elements control an elaborate vesicle transport process necessary for efficient exocytosis during anaphase and subsequent eggshell formation.

## Discussion

Many outer kinetochore proteins have been observed in linear elements since Nuf2^him-10^ localization was first reported in oocyte meiosis I in *C. elegans*^36^. Their function has remained enigmatic given that kinetochore proteins are primarily understood for their function at chromosomes, not in the cytoplasm. Linear elements share many properties with the fibrous corona, an expansion of the kinetochore that facilitates microtubule capture early in mitosis^66^. While linear elements have been observed primarily in oocytes, kinetochore proteins were previously observed in “flares” extending from chromosomes in mitotic cells under checkpoint activating conditions^67^. In addition, ROD-1 can form filaments in mitotic cells under certain conditions^41^. Our finding that linear elements interact with the ER and vesicles and are required for microtubule mediated vesicle movement is a novel function for these structures. Linear element function is both temporally and spatially distinct from kinetochores since they form in the cortical cytoplasm and interact with membranes prior to NEBD, when kinetochores first interact with spindle microtubules. These results may explain why some kinetochore proteins have interactions with membrane trafficking proteins. We hypothesize that outer kinetochore proteins have a dual role in coupling chromosomes and membranes to microtubule-based transport.

The role of linear elements in vesicle transport appears to be part of an elaborate process that globally organizes the oocyte cytoplasm. We define several distinct stages of ER and vesicle organization from prophase I arrest through anaphase I, when exocytosis occurs. Linear element formation occurs after the nucleus moves distally and the ER and vesicles are cortically displaced. Microtubules and kinesin activity are required for distal nuclear positioning^68,69^, and we find that microtubules regulate ER and vesicle distribution as well. In mouse oocytes, vesicle movement to the cortex is mediated by actin^70^, which could also be involved in *C. elegans*. The vesicle clustering event mediated by linear elements occurs just prior to NEBD. Interestingly, spindle checkpoint components function at the nuclear envelope to time NEBD ^71^, which could also regulate linear elements. Indeed, spindle checkpoint and nuclear pore components localize to linear elements^43^. Following the clustering event, a dynamic process of cytoplasmic streaming occurs, which has been observed in multiple cell types^72^. Previously, the ER and microtubules were shown to be required for streaming, which distributes cortical granules in the cortex for efficient exocytosis in anaphase I^65^. The role of linear elements during streaming is not clear, but they remain relatively stationary in the cortex while the ER, vesicles and microtubules flow with the cytoplasm. Interestingly, depletion of CLS-2, which is observed at linear elements, disrupts the microtubule cortex and affects polar body extrusion^45^. Finally, kinesin packs organelles inward to facilitate spindle positioning^68,73^. Depletion of the KCA-1 kinesin adapter was previously shown to inhibit cortical granule transport to the cortex, causing vesicles to be retained in meiosis II^29^. The fact that multiple mechanisms govern cytoplasmic organization in the oocyte may explain why depletion of linear element kinetochore proteins does not cause a severe vesicle exocytosis defect. Additionally, our RNAi conditions may not be penetrant enough or remaining kinetochore complexes have sufficient activity to move vesicles to their destination. Multiple mechanisms may be redundant or act in parallel, with linear elements serving to enhance the efficiency of the process. Future studies will be required to fully understand the role of linear elements during vesicle transport in the oocyte.

One important question is what mechanism(s) are required for linear element assembly. We observe small puncta rapidly increasing in size to intermediate and full-length structures. This pattern is reminiscent of cytoophidia, structures made of polymerized CTP synthase that start as small puncta that fuse into similar elongated filamentous structures^74^. This suggests a stepwise formation process with a hierarchy of assembly, as is the case for kinetochore formation on chromosomes^75^. Indeed, recruitment of KNL-1, CLS-2 and BUB-1 to linear elements exhibit hierarchical dependency^45^. We show that the small puncta labeled with CZW-1^ZW10^ do not enlarge after depletion of Nuf2^him-10^ at NEBD, indicating that Nuf2 is required for efficient linear element assembly. Purified RZZ-Spindly proteins can assemble into filaments in vitro^41^. RZZ-Spindly exhibits structural similarity to vesicle coat proteins^51^, which polymerize to organize donor membranes into vesicles. Therefore, RZZ-Spindly oligomerization may drive linear element formation as it does with fibrous corona expansion^66^. Future studies will be required to understand the assembly of linear elements based on outer kinetochore protein interactions.

In addition to the intrinsic ability of kinetochore proteins to polymerize, their interactions with microtubules also contribute to linear element formation. We observe that linear elements maintain a constant and dynamic colocalization with the cortical microtubule network. Furthermore, microtubule depletion inhibits linear element growth at the time of NEBD. Linear elements expand after treatment with nocodazole^41,49^, or when we deplete microtubules and inactivate APC/C. How microtubules affect the size of linear elements is unclear. The lack of linear element growth without Nuf2^him-10^ may reflect a requirement for plus end microtubule binding in linear element assembly. Collectively, our results suggest that the intrinsic ability of RZZ-Spindly to polymerize is enhanced by microtubule-dependent mechanisms *in vivo*. Multiple microtubule interacting proteins are observed at linear elements that may facilitate linear element formation. RZZ-spindly is well known for its role in recruiting dynein to kinetochores^76^, which might also be involved in linear element assembly. Our finding that Nuf2^him-10^ depletion causes formation of small and motile linear element puncta might reflect residual dynein-mediated movements. Future studies will be important to understand how kinetochore proteins and microtubules contribute to linear element formation.

Since linear elements form just prior to NEBD and disappear during anaphase I, we suspect that they may be under the regulatory control of cyclin dependent kinase, which is first activated in the cytoplasm^77^. The fibrous corona can be detached from chromosomes after microtubule and CDK-1 inhibition^41,66,78^. Previously, dynein was observed on linear element-like structures, which grow extremely large when APC/C, microtubules and CDK are inhibited^79^. Both APC/C and CDK-1 depletion cause eggshell defects in *C. elegans*, but the underlying mechanism(s) are mostly uncharacterized. Inhibition of numerous cell cycle genes was previously shown to affect the biogenesis, transport and exocytosis of cortical granule vesicles^33^. CDK-1 depletion inhibits cell cycle progression in meiosis I and prevents cortical granule exocytosis^33^. APC/C inhibition blocks cell cycle progression past prometaphase I and causes retention of cortical granules that accumulate in clusters^33^. APC/C and CDK-1 activity regulate motor activity to control spindle positioning and rotation during meiosis I and are required for cytoplasmic streaming^69,80,81^. Whether APC/C and CDK-1 affect cytoplasmic organization through linear elements in addition to their roles at the spindle is an important question for future studies. The interplay between microtubule dynamics, cell signaling and kinetochore protein polymerization likely control linear element behavior and function.

A critically important question is to understand how linear elements couple microtubules to vesicle transport. The simplest model suggested by kinetochore function is that linear elements would assemble on vesicles and bind to the plus end of microtubules to pull vesicles to their destination. However, linear elements initially form in close contact with the ER and microtubules and gradually associate with vesicles as they assemble into larger structures. The observation that ZW10 exists in an ER tethering complex supports a model where linear elements first form at the ER^52,53^. As linear elements assemble, they dynamically appear to search and probe the cytoplasm, dynamically interacting with vesicles as they form clusters. This behavior contrasts with kinetochores, which assemble on chromosomes and serve as binding platforms for spindle microtubules, which are dynamic and search the cytoplasm. Given that different kinetochore proteins can enable microtubule binding at the plus and minus ends as well as laterally^49,82^, each of these modalities may contribute to vesicle transport. While we find that microtubules facilitate linear element assembly, linear elements may also affect microtubule organization. Vesicles could be transported by motors such as dynein, well known for its role in vesicle transport^83^, that operate on the microtubule network, on linear elements, or both. Future studies using precise mutations that can uncouple linear element formation from microtubule and motor function will be required to tease out these mechanisms. In the future, studies of linear elements may reveal insights into vesicle transport mechanisms and serve as a useful context to further understand kinetochore protein function and behavior.

## Acknowledgements

*C. elegans* strains were provided by the CGC, based on information found in Wormbase, which are funded by the NIH (P40 OD010440) and NHGRI (U41 HG002223). We thank Dhanya Cheerambathur for sharing the KNL-1::TagRFP endogenously tagged strain. We thank Hayden Norris and Chris Turpin for data, strains and feedback. Funding was provided by the NIH grant R01 GM114471 to J.N. Bembenek.

---

**Figure S1.**
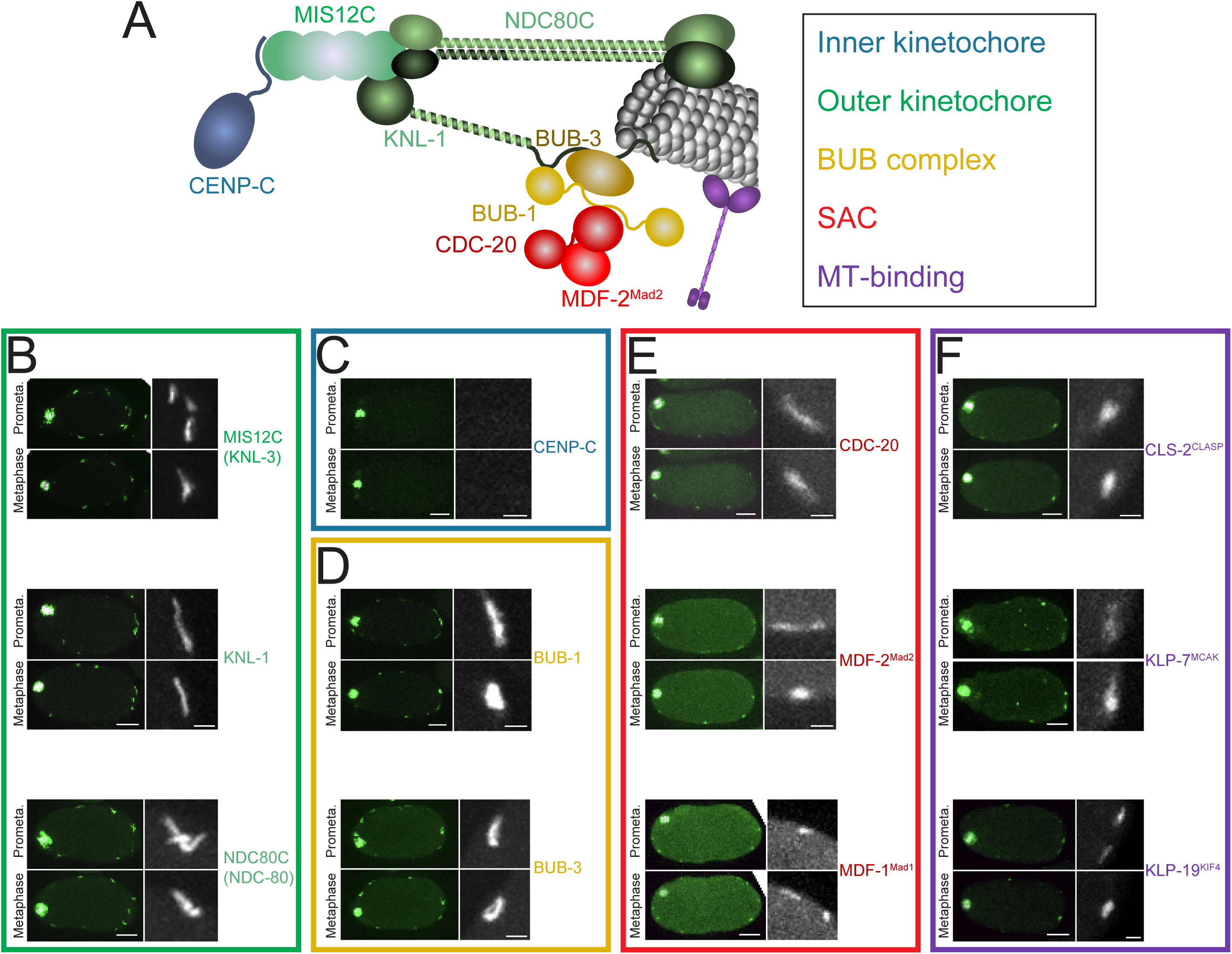
Analysis of Linear Elements Composition. (A) Schematic of the different proteins or protein complexes analyzed. (B-F) Prometaphase and Metaphase oocytes are shown for each component. A zoomed image of a single linear element is shown on the right of each oocyte. Scale bars, 10 µm (full oocytes) and 2 µm (zoomed images). (B) Components of the KMN network (C) Inner kinetochore (D) The BUB complex (E) SAC components that form the MCC (F) Microtubule-binding proteins

**Figure S2.**
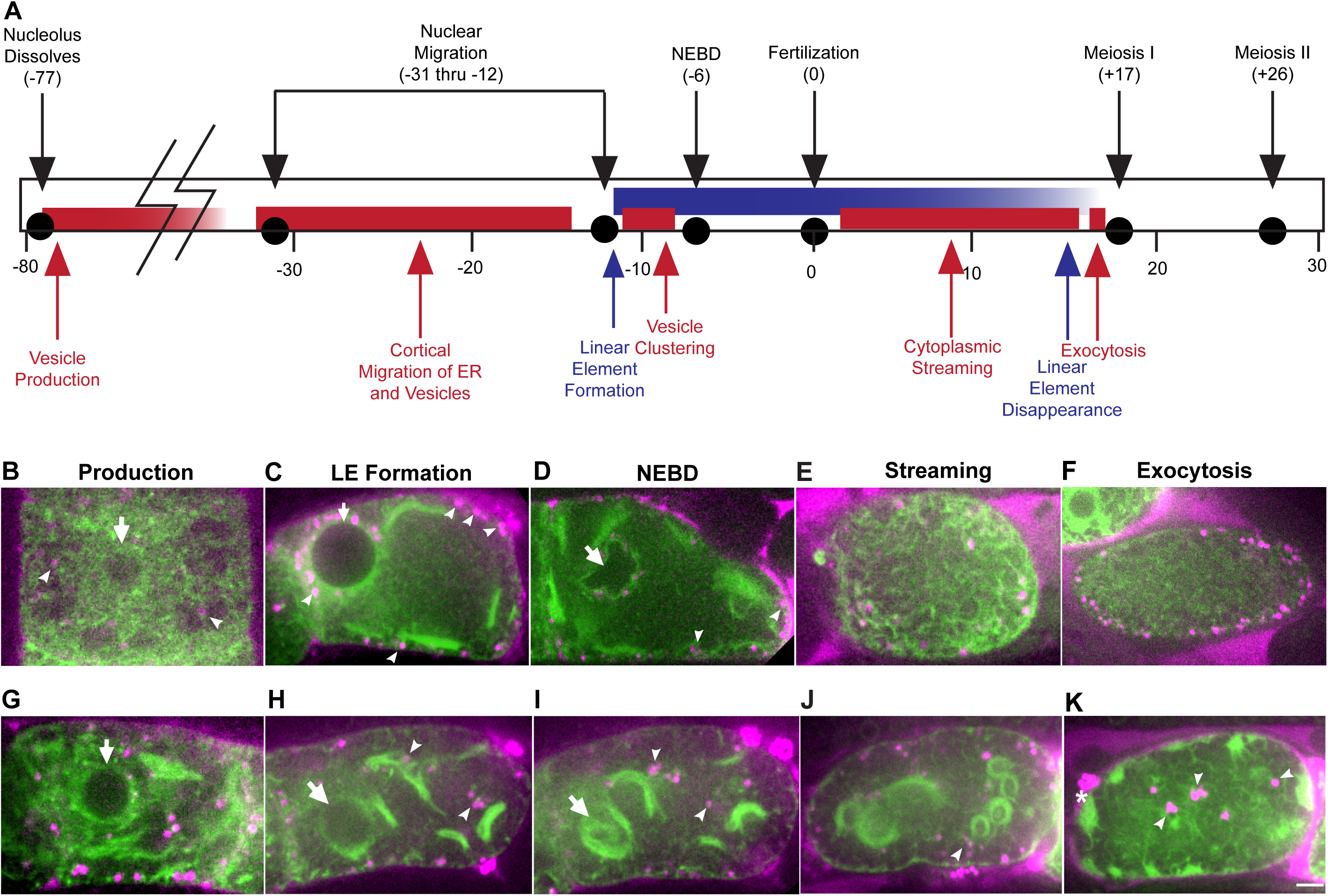
Global dynamics of ER and Cortical Granules during Meiosis. **I** (A) Timeline of events during oocyte meiosis. Upper events are based on McCarter et al^64^, events below are based on our observations of ER and vesicles during these different time periods. Red highlights vesicle events, blue indicates linear element timeline. (B-F) Representative images showing ER morphology and vesicle distribution at different stages of oocyte meiosis I. Arrows indicate position of the nucleus, which moves posteriorly by the time of filament formation. Arrowheads indicate cortical granules. (G-K) Organization of the ER and vesicles in the cytoplasm is significantly disrupted in *tba-2(RNAi)* oocytes at each stage of meiosis I. Arrows indicate nucleus, arrowheads indicate vesicles that are not cortically displaced in (H-J) and remain in the cytoplasm after they should normally undergo exocytosis in (K).

**Figure S3.**
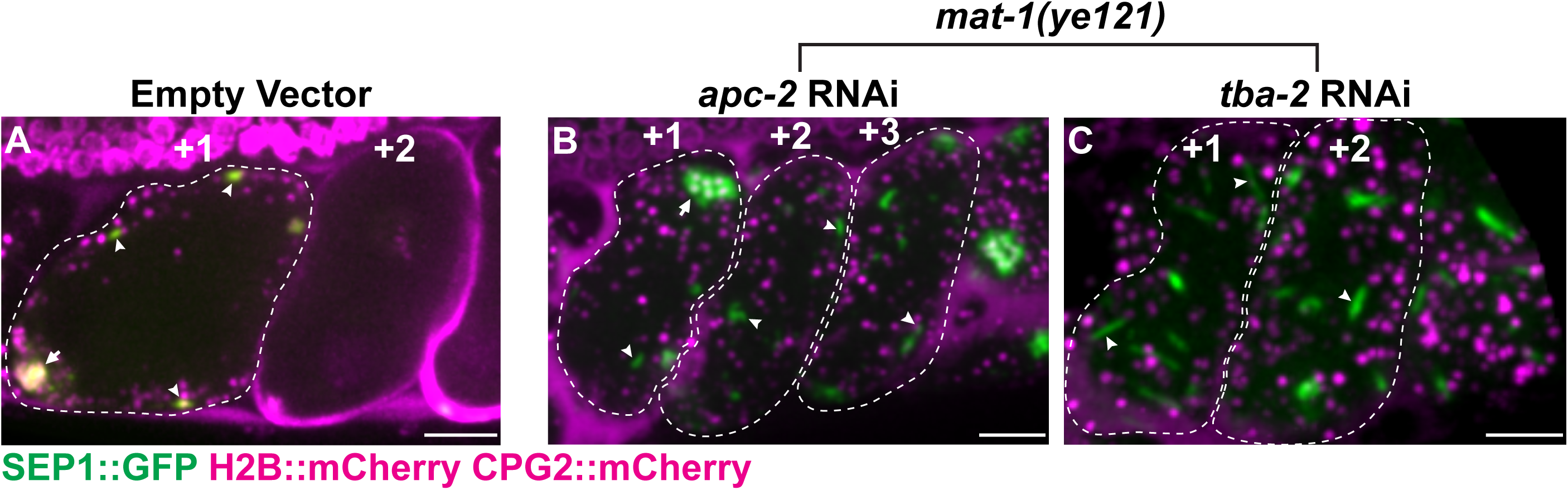
Linear Elements in APC/C Arrested Embryos. (A-C) Maximum Z projections of embryos expressing SEP-1::GFP to show linear elements, H2B::mCherry to label chromosomes, and CPG-2::mCherry in cortical granules. Scale bars are 5 µm. (A) Control embryos fed empty vector RNAi have numerous cortical granules throughout the cortex with multiple linear elements (arrowheads) during prometaphase I (^+^1 embryo, indicated by dashed outline). +2 and older embryos show CPG-2::mCherry incorporated into the eggshell after exocytosis and linear elements are not observed. (B) APC/C temperature sensitive mutant *mat-1(ye121)* embryos arrested in prometaphase I retain cortical granules in the cytoplasm and numerous linear elements (arrowheads) are observed. (C) *mat-1(ye121); tba-2(RNAi)* embryos arrested in prometaphase I also do not secrete CPG-2::mCherry. Despite microtubules affecting linear element formation at NEBD, long term arrested embryos have numerous linear elements (arrowheads) that appear longer than normal.

**Figure S4.**
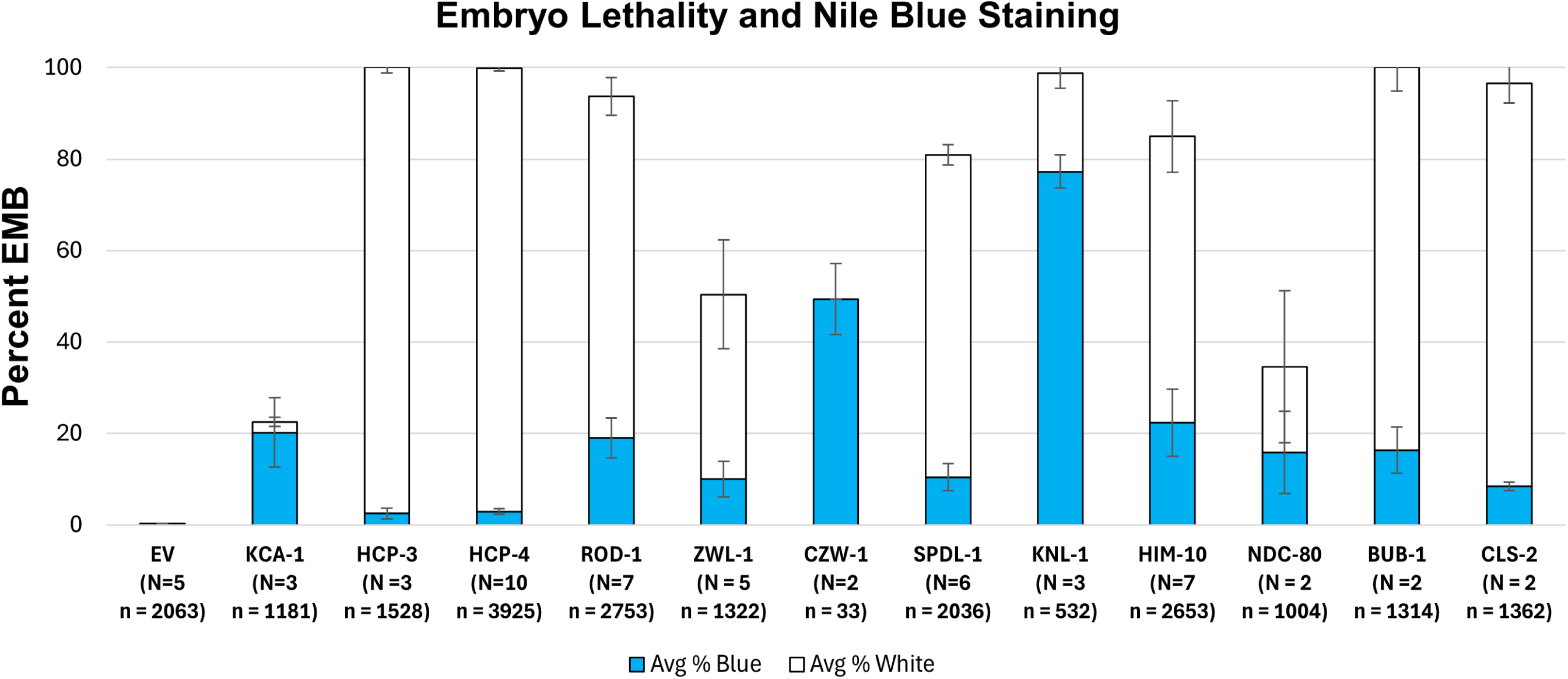
Depletion of Kinetochore Proteins Causes Eggshell Permeability. N2 animals were treated with the indicated feeding RNAi for 24-48 hours and the second day broods were assessed for eggshell permeability by nile blue A plate-based staining. Bars indicate total lethality, blue indicates the portion of dead embryos that show nile blue A staining. Depletion of the kinesin microtubule motor adapter, KCA-1, causes significant permeability among dead embryos as previously shown^84^. RNAi of *czw-1* is already known to cause eggshell permeability and causes low brood size on day 2, indicating a severe defect. Several other outer kinetochore proteins cause a low rate of eggshell permeability, more than depletion of centromeric (CENP-A^hcp-3^) and inner kinetochore (CENP-C^hcp-4^) proteins. Error bars indicate SEM. N indicates the number of experimental replicates for each condition, n is number of embryos.

## Materials and Methods

### Maintenance of Transgenic Worm Strains

Multiple *C. elegans* strains were isolated and maintained using traditional methods per Wormbook Protocols. Unless otherwise noted, worms were maintained at 20°C and fed E. Coli (OP50) supplied by the CGC. Some strains were obtained from the Caenorhabditis Genetics Center. Transgenic lines generated for this project are listed below in Table S1.

### RNAi Treatments

Feeding RNAi was conducted using bacterial strains from the Ahringer library. For feeding RNAi, L4 hermaphrodites were plated onto 60 mm agar NGM plates seeded with IPTG-induced RNAi bacterial strains at 20°C. Resultant phenotypes were assessed after 20-48hrs of treatment.

#### Nile Blue Staining

The permeability of embryos was assessed via protocol previously described^84^. 150 µg/ml Nile Blue A stain was added to the media prior to pouring. Plates were seeded with HT115 bacteria harboring the RNAi construct as described above. L4 worms were fed various RNAi strains over the course of 24 hours and allowed to lay embryos. Animals were removed and placed onto fresh RNAi seeded plates containing Nile Blue A and allowed to lay embryos for another 24 hours until removal. Laid embryos were left on the plate for 24 hours after parent removal and were assessed for blue staining and embryo lethality using a SMZ745 Nikon Stereomicroscope under white light. Embryos containing intact eggshells were impermeable to the dye, and embryos with defective eggshells were stained blue. Stained embryos, unstained embryos, and hatched larvae are expressed as a percentage of the total brood, averaged among 2-10 independent experiments.

### Live cell imaging

Live cell imaging data of JAB19, JAB274, JAB276, JAB280, JAB285, JAB286, DKC547 were collected on an inverted Nikon Eclipse Ti2-E with a 60 X 1.42NA objective and 100 X 1.45 NA objective, a CSU-X1 spinning disk system, and Andor iXon Life camera operated by NIS-Elements software (Nikon). Unless otherwise mentioned, live cell imaging was conducted at room temperature. Live imaging of KLP-19 (FGP36), CDC-20 (OD2591), and KNL-3 (FGP722) was done using a CFI Plan Apochromat Lambda 60×/NA 1.4 oil objective mounted on a microscope (Nikon Eclipse Ti) equipped with a Prime 95B 22 mm camera (Photometrics), a spinning-disk head (CSU-X1; Yokogawa Electric Corporation).

Acquisition parameters were controlled with NIS software (Nikon).

Live imaging of HCP-4 (FGP311), NDC-80 (FGP372), KNL-1 (FGP517), MDF-1 (OD2920), MDF-2 (OD216), BUB-1 (FGP202), BUB-3 (TG4193), KLP-7 (FGP261), CLS-2 (JDU38) was done using a 60×/NA 1.4 oil objective mounted on a microscope (IX81; Olympus), an EMCCD Cascade II camera (Photometrics), spinning-disk head (CSU-X1; Yokogawa Electric Corporation). Acquisition parameters were controlled by MetaMorph seven software (Molecular Devices).

For all live imaging experiments, full or partial projections, and single planes are presented. Files were stored, classified, and managed using OMERO (Allan et al., 2012). Figures were prepared using OMERO.figure and assembled using Adobe Illustrator. Image analysis and manipulation was performed in Fiji (National Institutes of Health).

#### In Utero Live Cell Imaging

A chemical immobilization method was applied by mounting worms in an M9 solution containing 5mM levamisole (2% solution) on a 2% agarose pad affixed to a glass slide and sealed with Vaseline, following standard protocol^33^.

#### Ex Utero Time Lapse Imaging of Meiosis I

Oocytes were dissected and mounted in 5 µl of L-15 blastomere culture medium (0.5 mg/mL inulin; 25 mM HEPES, pH 7.5 in 60% Leibowitz L-15 medium and 20% heat-inactivated FBS) on 24 × 40 mm #1.5 coverslips. Once dissection was performed and early oocytes were identified using a stereomicroscope, a circle of Vaseline was laid around the sample, and a custom-made plastic holder (with a centered window) was placed on top of the coverslip. The sample was imaged immediately.

### Quantifications

#### Fluorescence

Fluorescent values for the linear element and chromosome vs. cytoplasm curve in Fig. 1 were acquired from single plane, *in utero* movies of maturing –1 oocytes using the same acquisition settings. The values represent the individual signal from independent movies, found in a 3-pixel diameter circle at the linear elements, and chromosomes. This was repeated using a 15-pixel diameter circle for cytoplasm and background signals. The values for each timepoint at these regions of interest are averages of three independent measurements, minus the average background signal.

#### Statistics

For the Nile Blue data, brood size was calculated as the sum of the embryos and larvae present on a 35mm NGM plate 48hrs after feeding. For each RNAi condition, the percentages for white/blue embryos as a fraction of brood size was averaged across 2-10 experiments and plotted as a stacked column. Error bars display the standard error of the mean (S.E.M) for white/blue embryos for each condition.

---

**Table S1.**
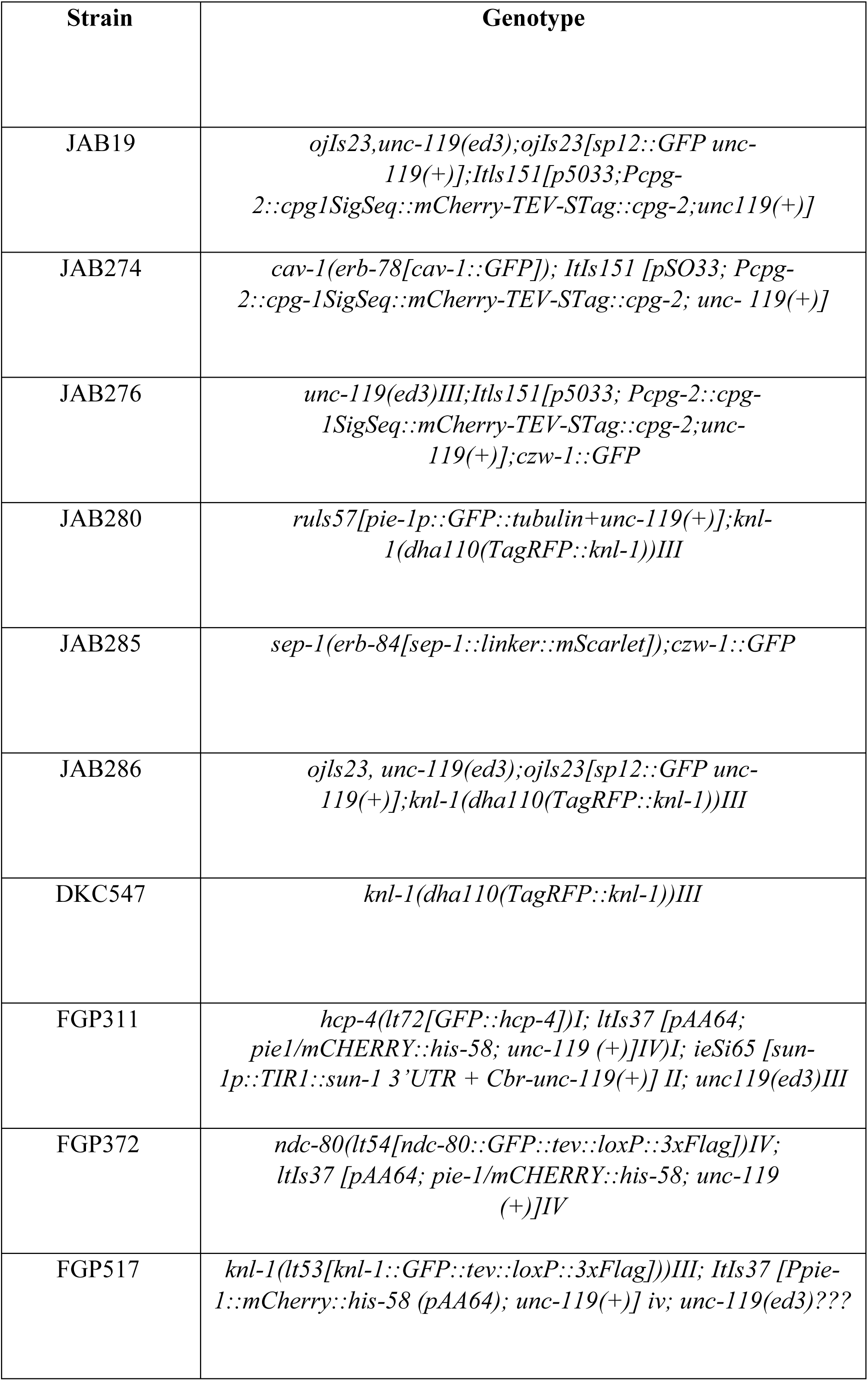
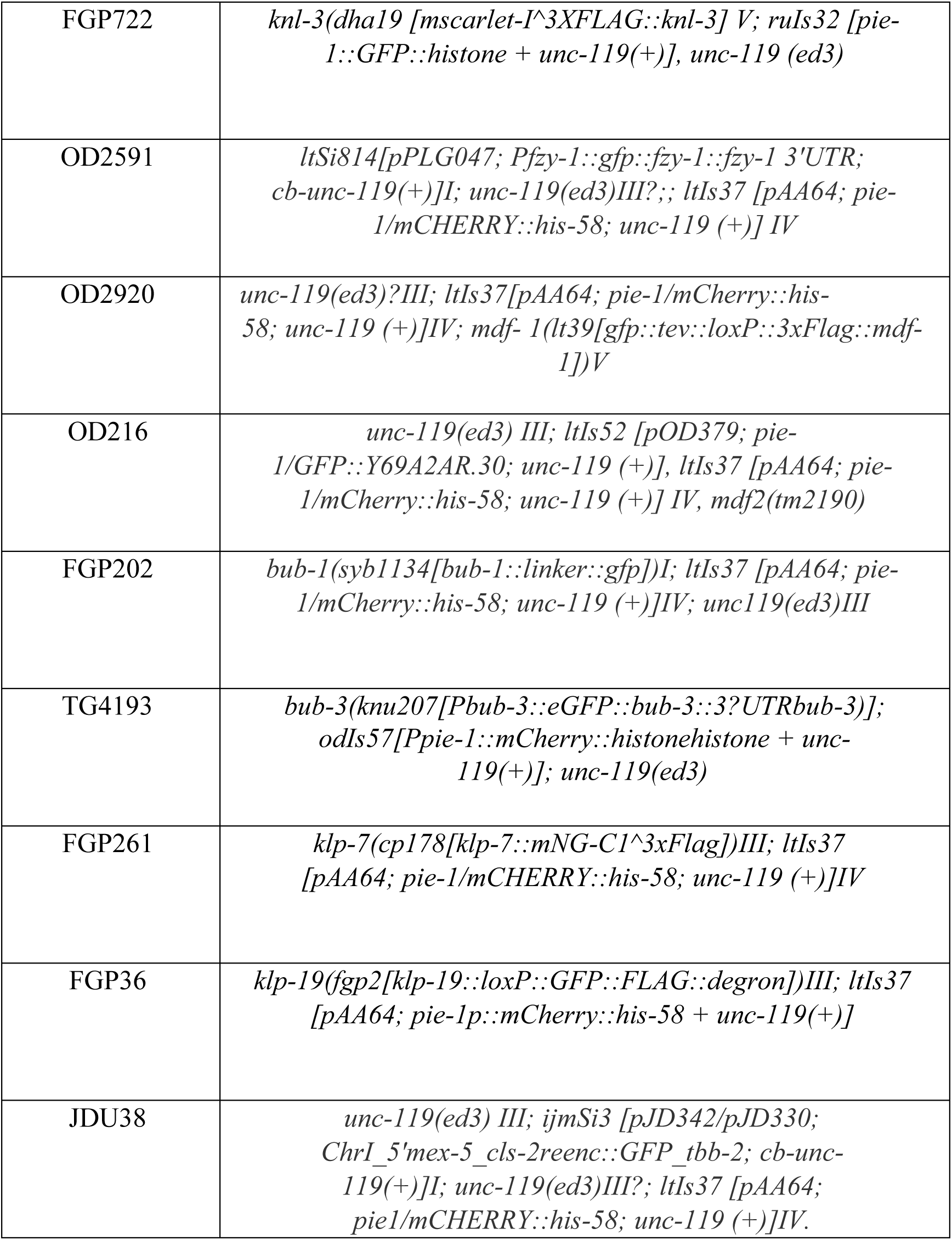
List of *C. elegans* strains used in this study.

